# Modelling the emergence of an egalitarian society in the n-player game framework

**DOI:** 10.1101/305912

**Authors:** Kohei Tamura, Hiroki Takikawa

## Abstract

Unlike other primates, human foragers have an egalitarian society. Therefore, the evolution of egalitarian behaviour has been the subject of long-standing debate in a wide variety of disciplines. A recent hypothesis states that a social control against potentially dominant individuals played an important role in the emergence of an egalitarian society, although this has not been modelled directly. In the present study, we modelled this hypothesis based on the n-player game framework, in which the owner, who may attempt to monopolise resources, could be punished by a coalition of other group members. Our results suggest that a potentially despotic payoff structure can promote the evolution of egalitarian behaviour. Besides, large group size, small cost of competition, and variation in the strengths of individuals can promote the evolution of egalitarian behaviour. Our results suggest the importance of both social control against dominant individuals and benefits of a coalition for the evolution of egalitarian behaviour.

## 1 Introduction

The emergence of an egalitarian society among foragers has been a long-standing question in biology, anthropology, and other social sciences. Apes, which are phylogenetically close to humans, are known for forming rather strong dominant hierarchies with authoritarian leadership [1,2]. Although humans have also developed highly unequal social systems ranging from chiefdoms to the modern states [3], psychological and economic studies have reported a large body of evidence supporting egalitarian motives of humans in modern industrialised societies [4–6].

The key to understanding the development of an egalitarian society is egalitarian ethos. Ethnographic studies reported the existence of egalitarian ethos in a wide variety of forager societies [7]. In addition, the innate human tendency of aversion to inequality has been examined by recent behavioural experiment studies [6,8–12]. Thus, the hypothesis that claims that the evolution of egalitarian ethos enabled humans to develop and maintain an equal social system is worth extensive investigation.

The theory of the evolution of egalitarian ethos was mostly elaborated in Boehm’s series of seminal works [13–16]. Based on extensive evidence from Late-Pleistocene-appropriate foraging societies, he hypothesised that social control plays a central role in the evolution of egalitarian ethos. Foragers have a social control system to punish those who violate egalitarian norms or use bossy behaviours. For example, if a hunter, who is strong and has potential to become an alpha male, tried to monopolise game meat, other members would form a coalition and collectively punish the norm-violating member. This greatly reduces the fitness of would-be alpha males, thus leading to the evolution of egalitarian ethos and norm-confirming behaviours.

The remaining question to be addressed is, ‘Which conditions exactly allow collective punishment to be effective and promote the evolution of egalitarian ethos?’ To answer this question, mathematical modelling of the evolution of egalitarian ethos is employed.

A number of mathematical models of the evolution of egalitarian behaviour have been developed based on various frameworks including the ultimatum game [17–20], bargaining game [21], prisoner’s dilemma with punishment [22,23], and hawk-dove game. Among them, Gavrilets’s [24] model is the one most directly related to collective punishment for monopolisers, as it models owner-bully-helper interactions. The ‘owner’ has an item, and the ‘bully’ may try to take it from the owner. The ‘helper’ may decide to form a coalition with the owner to fight the bully. Using numerical simulations, he showed that, under some conditions, the evolution of helping behaviour can occur. The key assumption here is that a fitness function takes the form of a generalisation of the Tullock contest success function [25]. This implies that stopping the bully may contribute to improvement of the fitness of not only owners but also helpers.

Gavrilets’s model provides several useful insights. Among others, employing a generalisation of the Tullock contest success function as a fitness function is crucial to understanding the evolution of an egalitarian society from a game-theoretical viewpoint.

However, the assumption of Gavrilets’s [24] model seems to be different from the ethnographic observations and Boehm’s hypothesis in some respects. First, it assumes that, maximally, two individuals attempt to punish the norm violator. However, ethnographic reports suggest that all group members punished the norm violator; therefore, the n-player game framework, in which all group members make their decision, seems more appropriate. Second, Boehm’s hypothesis assumes that the owner of the resource would be punished if he or she would not share the resource; however, in Gavrilets’s model, the bully who attempted to take the resource from the owner is the individual to be punished. In the present study, to model Boehm’s hypothesis more directly, we use the n-player game framework. We investigate which factors can affect the evolution of egalitarian behaviour, or the coevolution of resource sharing and punishment of monopolisers.

## 2 Model

We consider a population of infinite individuals with non-overlapping generations. For simplicity, we assume that these individuals reproduce asexually. At the beginning of each generation, individuals form a group consisting of *n* individuals. In each group, an individual is randomly chosen and becomes the owner of an amount of resource, *R*. We refer to an individual who is not an owner as a peer. The owner equally allocates the resource within the group or attempts to monopolise it. The individuals are divided into two types, *E*_0_ and *E*_1_, based on their behaviour as an owner. On one hand, an *E*_0_ owner attempts to monopolise the resource, which may lead to competition within the group. On the other hand, an *E*_1_ owner shares the resource so that all group members, including the owner, obtain *R/n*. The individuals are also classified into two types, *C*_0_ and *C*_1_, according to their behaviour as a peer. A *C*_0_ peer does nothing against the monopolisation by the owner. If the owner shares the resource, a *C*_1_ peer does not form a coalition and enjoys the resource *R/n*, whereas, if the owner attempts to monopolise the resource, *C*_1_ peers form a coalition to penalise the owner. If there is at least one *C*_1_ peer, competition over the resource occurs. We use the Bradley–Terry model to describe the probability of individual (or group) *i* winning against individual (or group) *j*, which is defined by

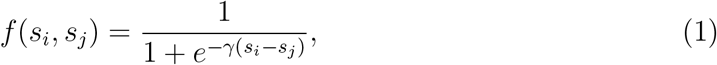

where *s_a_* is the strength of an individual or coalition *α*, and *γ* is a parameter to regulate how difference in strength affects the result of the competition. Individuals are dichotomised into two categories based on their strength— strong or weak—and their strength is denoted by *s_s_* and *s_w_*, respectively. We assume that the strength of individuals is determined during their developmental process: an individual becomes strong or weak with probabilities *ϕ* and 1 – *ϕ*, respectively. Let *k_s_* and *k_w_* denote the numbers of strong and weak individuals in a coalition, respectively. The strength of the coalition of *k* = (*k_s_* + *k_w_*) individuals, *S*(*k_s_*,*k_w_*), is defined by 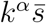
, where *k* is the number of individuals who join the coalition, 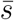 is the average strength of the coalition (= [*k_s_s_s_* + *k_w_s_w_*]/*k*), and *α* (> 0) is a parameter to regulate the synergy of the coalition. If the owner wins the competition, the owner gains resource *R*, and individuals who joined the coalition suffer the cost *c*. If the coalition wins, the owner suffers the cost *c*, and members of the coalition share the resource so that they gain an equal amount of the resource *R/k*.

There are four strategies, *E*_0_*C*_0_, *E*_0_*C*_1_, *E*_1_*C*_0_, and *E*_1_*C*_1_, which we refer to as ‘monopoliser’, ‘greedy’, ‘pacifist’, and ‘egalitarian’, respectively. Their frequencies are denoted by *x*_00_, *x*_01_, *x*_10_, and *x*_11_, respectively.

Let *π* and *ρ* denote the payoffs when the focal individual is the owner and peer, respectively. The probability that the numbers of strong and weak *C*_1_ individuals in a group of *n* are *k_s_* and *k_w_*, respectively; then *q*(*n*,*k_s_, k_w_*) is defined by

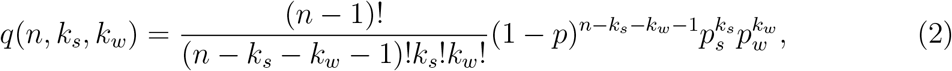

where *p* = *x*_01_ + *x*_11_, *p_s_* = *ϕp*, and *p_w_* = (1 – *ϕ*)*p*.

The payoffs of the four strategies as an owner are

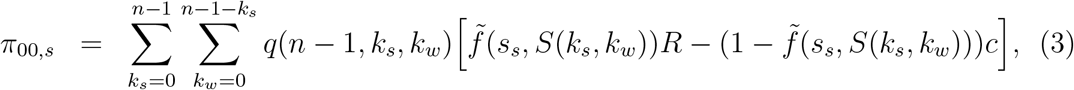

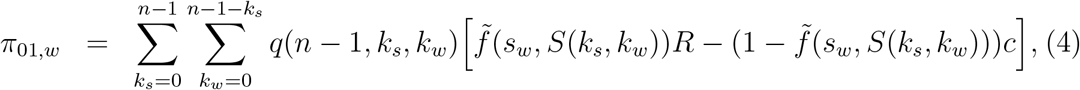

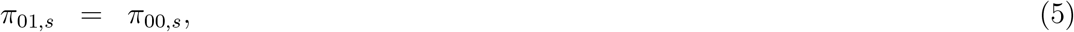

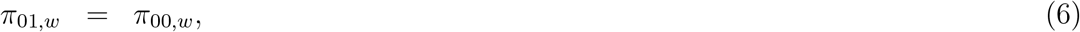

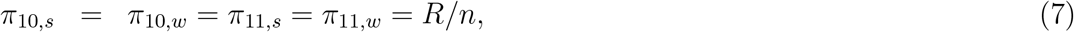

where

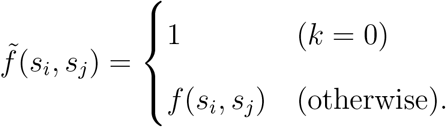

Because their behaviour as owners is identical, the payoffs of monopoliser and greedy individuals are the same depending on the strength of individuals. Further, the payoffs of pacifist and egalitarian individuals are identical irrespective of their strengths, because they share the resource.

Likewise, the payoffs of the four strategies as a peer are

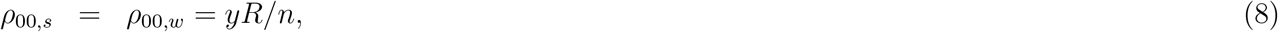

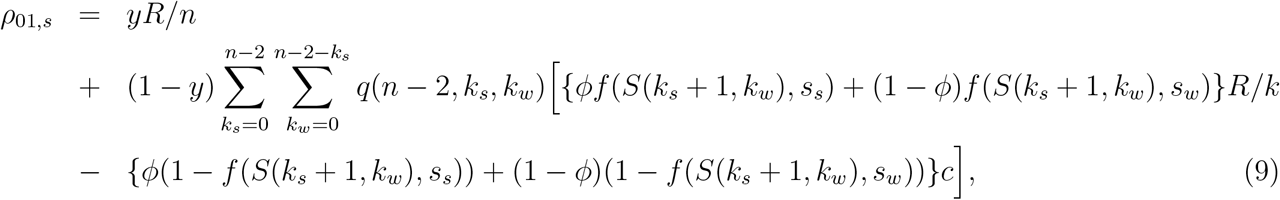

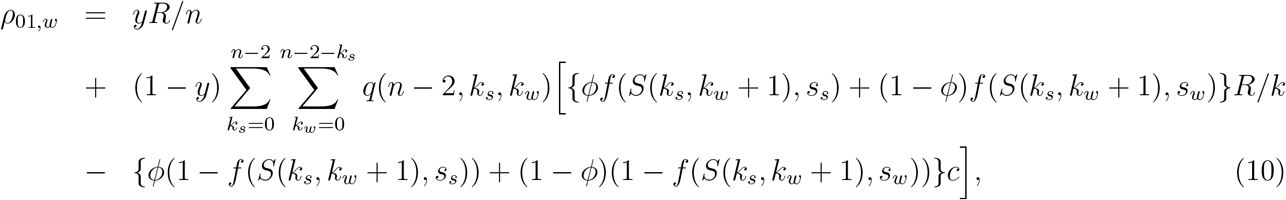

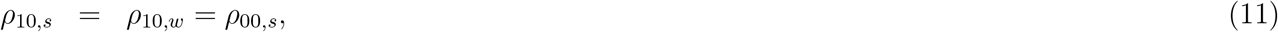

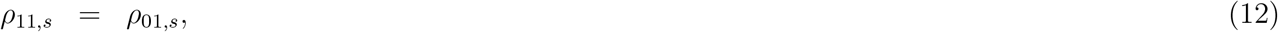

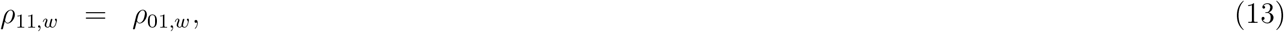

where *y* = *x*_10_ + *x*_11_. The payoffs of monopoliser and pacifist individuals are identical irrespective of their strengths, because they do nothing as a peer. Because their behaviour as peers is identical, the payoffs of greedy and egalitarian individuals are the same depending on the strengths of individuals.

The recursion equation is defined by

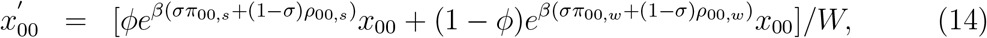

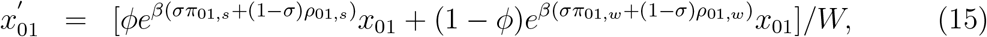

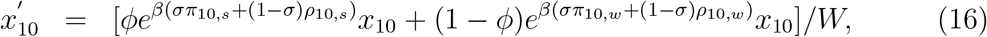

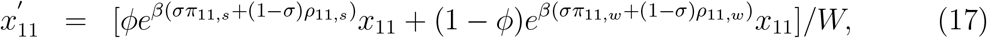

where *σ* represents the intensity of selection. For simplicity, we assume *σ* =1/2. *W* is the mean fitness of the population. *β* regulates the potential of inequality (0 ≤ *β*). A low value of *β* indicates a society wherein all individuals can enjoy the same reproductive success. Suppose, for example, when *β* = 0, all individuals receive the same amount of payoffs irrespective of their behaviour. A high value of *β* indicates the potential for disparity because a minor difference in the payoffs can be translated into a large difference in fitness.

Figure 1 shows the payoffs of the owner and peers in different situations.

## Numerical Analysis

We set *x*_00_ = 0.97, *x*_01_ = 0.01, *x*_10_ = 0.01, and *x*_11_ = 0.01 as the initial conditions. We regard the frequencies after 30,000 generations as equilibrium frequencies.

In what follows, we regard the egalitarian behaviour and an egalitarian society as a set of behaviours involving resource sharing and penalizing a norm violator and a population mostly occupied by egalitarian (*E*_1_*C*_1_) individuals, respectively.

## Results

To investigate which factors can promote the evolution of egalitarian behaviour, we consider as the initial condition that the population is almost occupied by monopolisers and examine a few other strategies that can invade the population.

**Figure 1:**
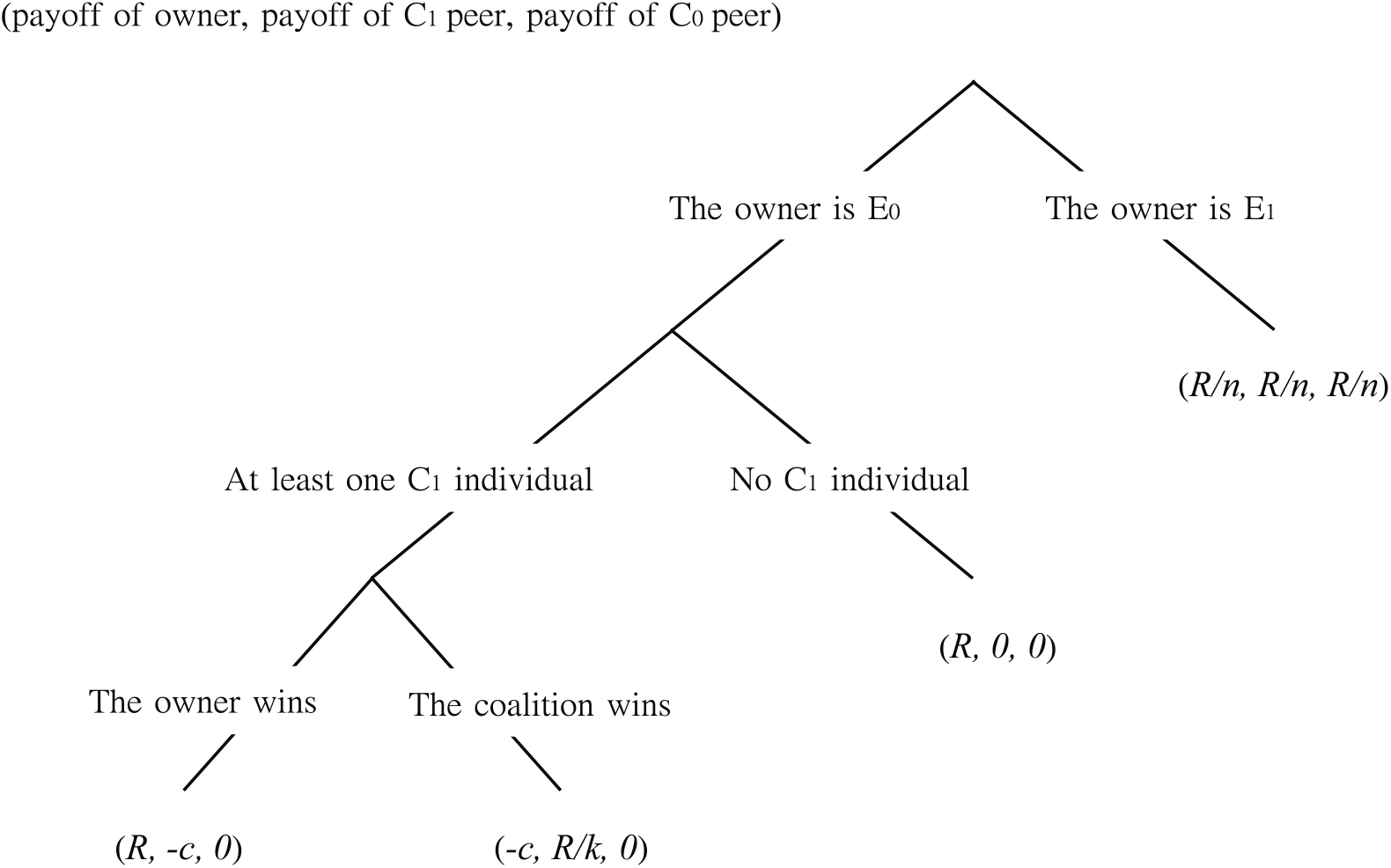
Payoffs of the owner and peers who join and do not join the coalition in different situations.

Figure 2(a) shows an example of the trajectory of the four strategies. First, the frequency of greedy individuals, who attempt to monopolise the resource when they are an owner and also form a coalition when they are a peer, increased. After the frequency of greedy individuals reached a certain level, the frequency of egalitarian individuals suddenly increased. The frequency of pacifist individuals remained low, although they did not become extinct. It should be noted that, once resource sharing behaviour was fixed in a population (i.e. the population was occupied by pacifist and/or egalitarian individuals), the frequencies did not change. This is because all individuals share their resources so that norm violators are not punished.

**Figure 2:**
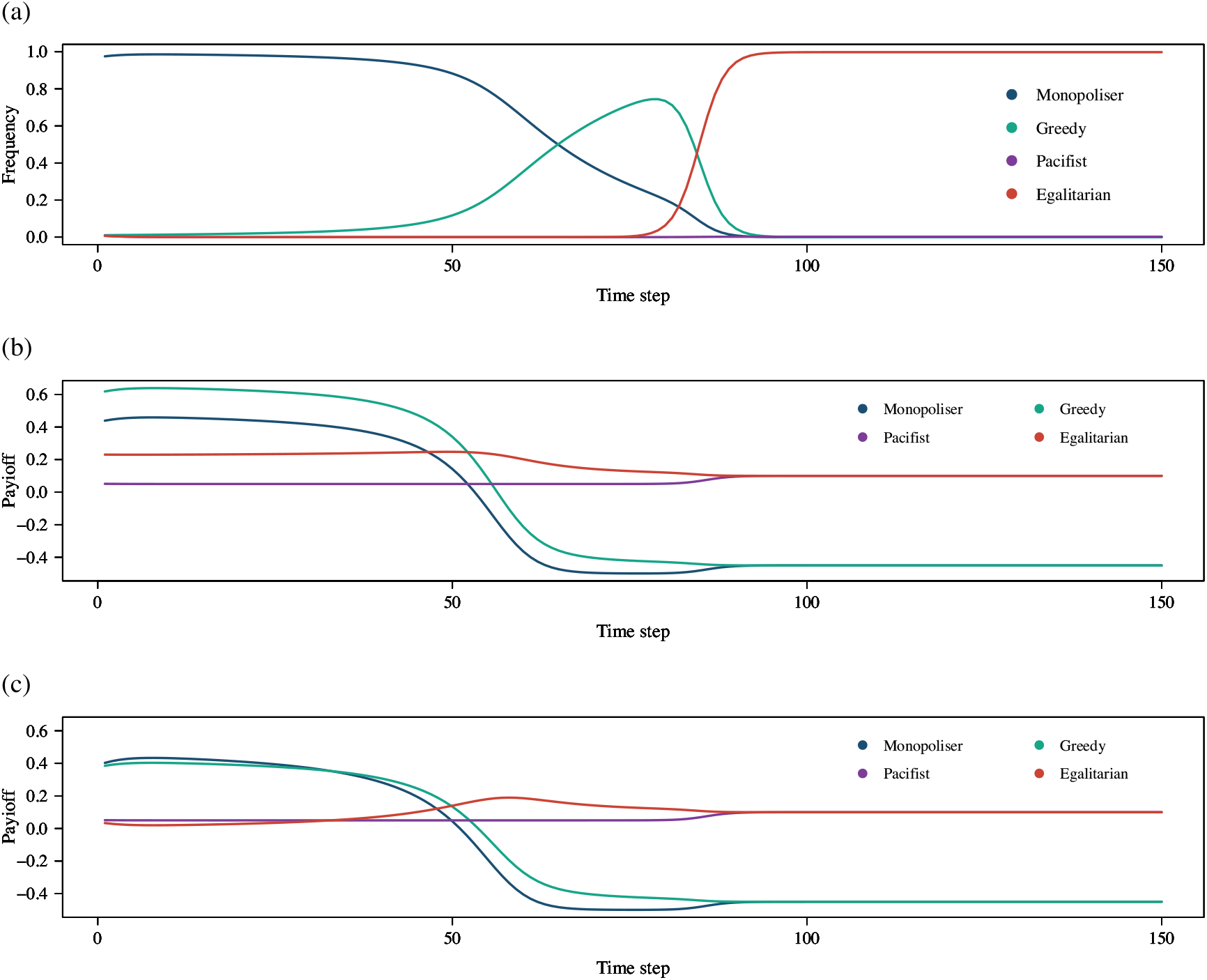
An example of changes in frequency and fitness of the four strategies. The initial frequencies of the four strategies are (*x*_00_,*x*_01_,*x*_10_,*x*_11_) = (0.97, 0.01, 0.01, 0.01). We set *γ* =1, *s_s_* = 2, *s_w_* = 1, *R* =1, *α* = 2, *β* = 1, *ϕ* = 0.25, *c* =1 and *n* = 10. (a) Frequency of the four strategies. (b) Fitness values of strong individuals. (c) Fitness values of weak individuals.

Figures 2(b) and (c) show an example of changes in the fitness of the four strategies. When the frequency of monopoliser individuals was large, the payoffs of monopoliser and greedy individuals were larger than those of other two strategies. For a strong individual, the payoff of a greedy individual was larger than that of a monopoliser individual (Figure 2 (b)). On the other hand, for a weak individual, the opposite was true (Figure 2 (c)). As the frequency of greedy individuals increased, the payoffs of monopoliser and greedy individuals decreased. This could be because, as the frequency of individuals forming a coalition (i.e. greedy and egalitarian individuals) increases, monopoliser and greedy individuals are more likely to be punished. The fitness of egalitarian individuals increased at first, associated with the decrease in the fitness values of monopoliser and greedy individuals, although the increase in fitness is larger for weak individuals than for strong individuals. This could be because weak individuals could win due to a larger number of group members joining a coalition. The fitness of egalitarian individuals also decreased when the fitness values of monopoliser and greedy individuals were below a certain level. The reason is as follows. First, when the frequency of monopoliser and greedy individuals is large, the coalition of egalitarian individuals can enjoy larger payoffs by taking resources from the owner rather than monopoliser and pacifist individuals. Second, after the frequency of monopoliser and greedy individuals decreases, punishment is hard to occur, indicating that egalitarian individuals can only receive equally distributed payoffs. The fitness values of strong and weak pacifist individuals were almost stable, although they became the same after the extinction of monopoliser and greedy individuals.

We also investigated which factors can affect the evolution of egalitarian behaviour. Figure 3 suggests that high values of *β*, small cost of competition, *c*, and a variation in the strength of individuals (i.e. an intermediate value of *ϕ*) can promote the evolution of egalitarian behaviour. Figure 4 also shows that large group size, *n*, can promote the evolution of egalitarian behaviour.

**Figure 3:**
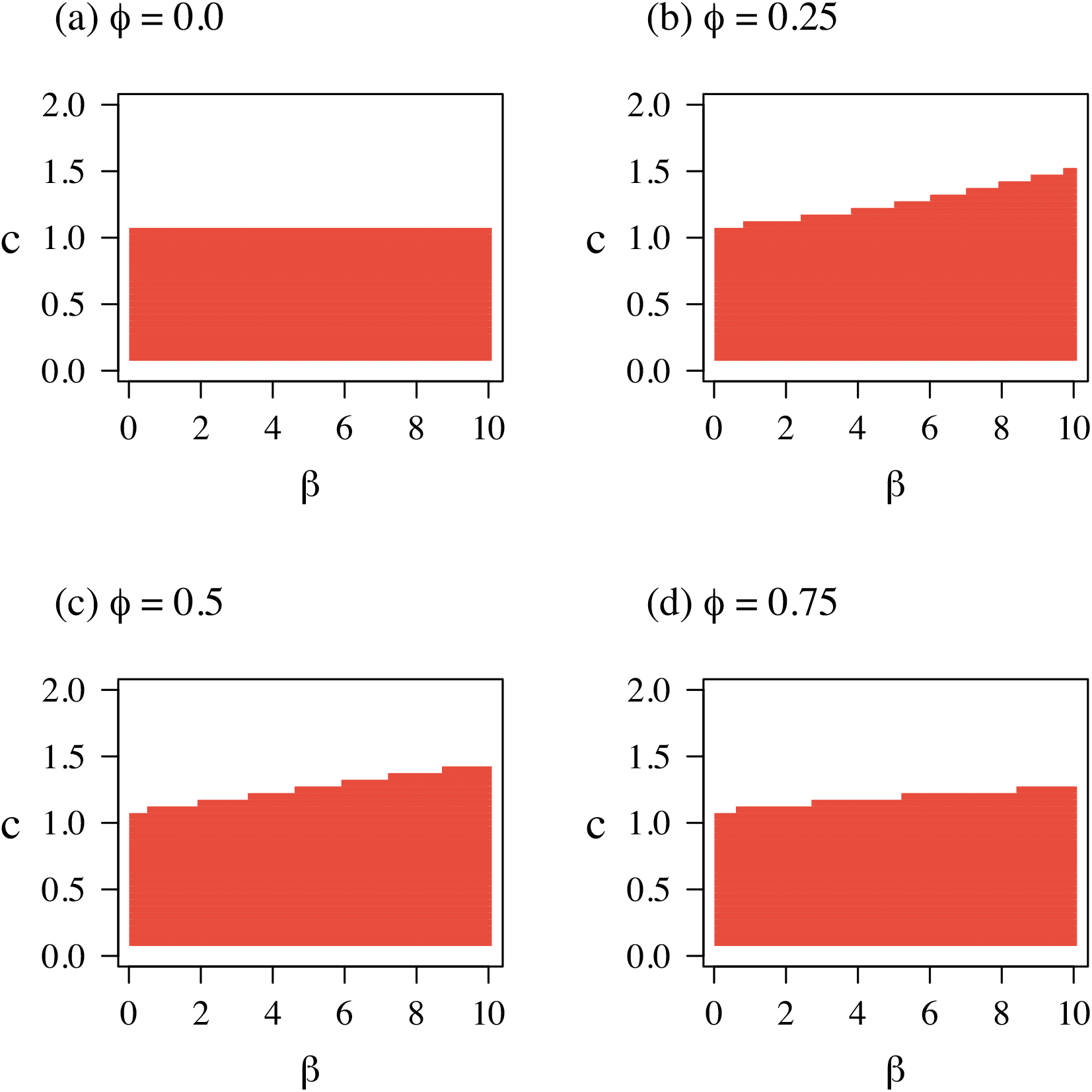
Evolution of egalitarian behaviour under various combinations of *β* and *c*. Red regions represent the fixation of *E*_1_ behaviour; that is, the equilibrium frequency of *y* is equal to unity. In addition, the frequencies of egalitarian individuals are close to unity in these regions. The initial frequencies of the four strategies are (*x*_00_,*x*_01_,*x*_10_,*x*_11_) = (0.97,0.01, 0.01, 0.01). We set *γ* =1, *s_s_* = 2, *s_w_* = 1, *R* =1, *α* = 2 and *n* = 10.

**Figure 4:**
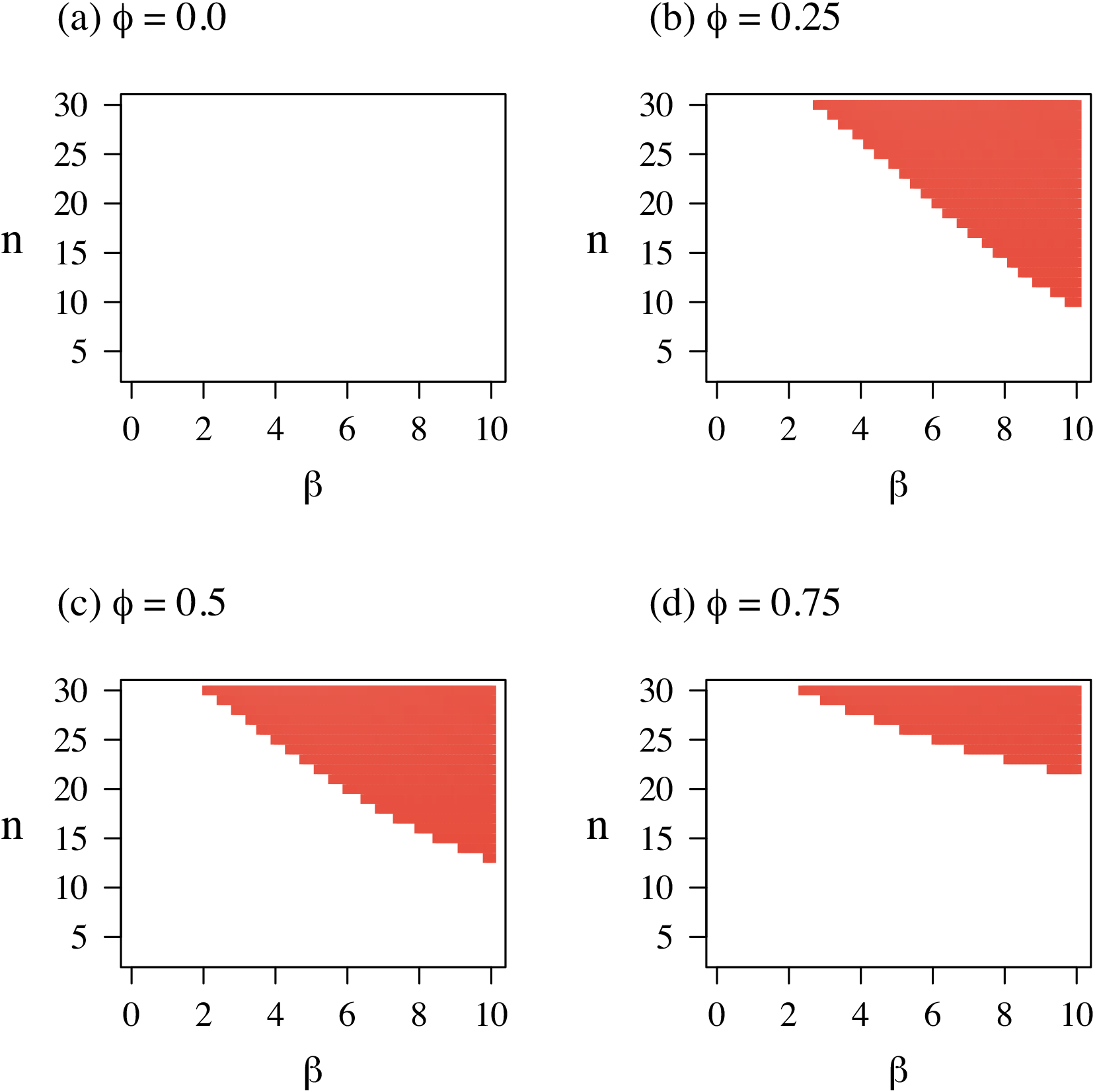
Evolution of egalitarian behaviour under various combinations of *β* and *n*. Red regions represent the fixation of *E*_1_ behaviour. The frequencies of egalitarian individuals are close to unity in these regions. The initial frequencies of the four strategies are (*x*_00_,*x*_01_,*x*_10_,*x*_11_) = (0.97, 0.01, 0.01, 0.01). We set *γ* =1, *s_s_* = 2, *s_w_* = 1, *R* =1, *α* = 2 and *c* = 1.5.

We further investigate the effects of group size, *n*, and *ϕ* on the probability of winning of the coalition and fitness of greedy individuals at the initial condition. Figure 5 shows effects of *n* on the probability of winning the coalition and the fitness value of greedy individuals.

**Figure 5:**
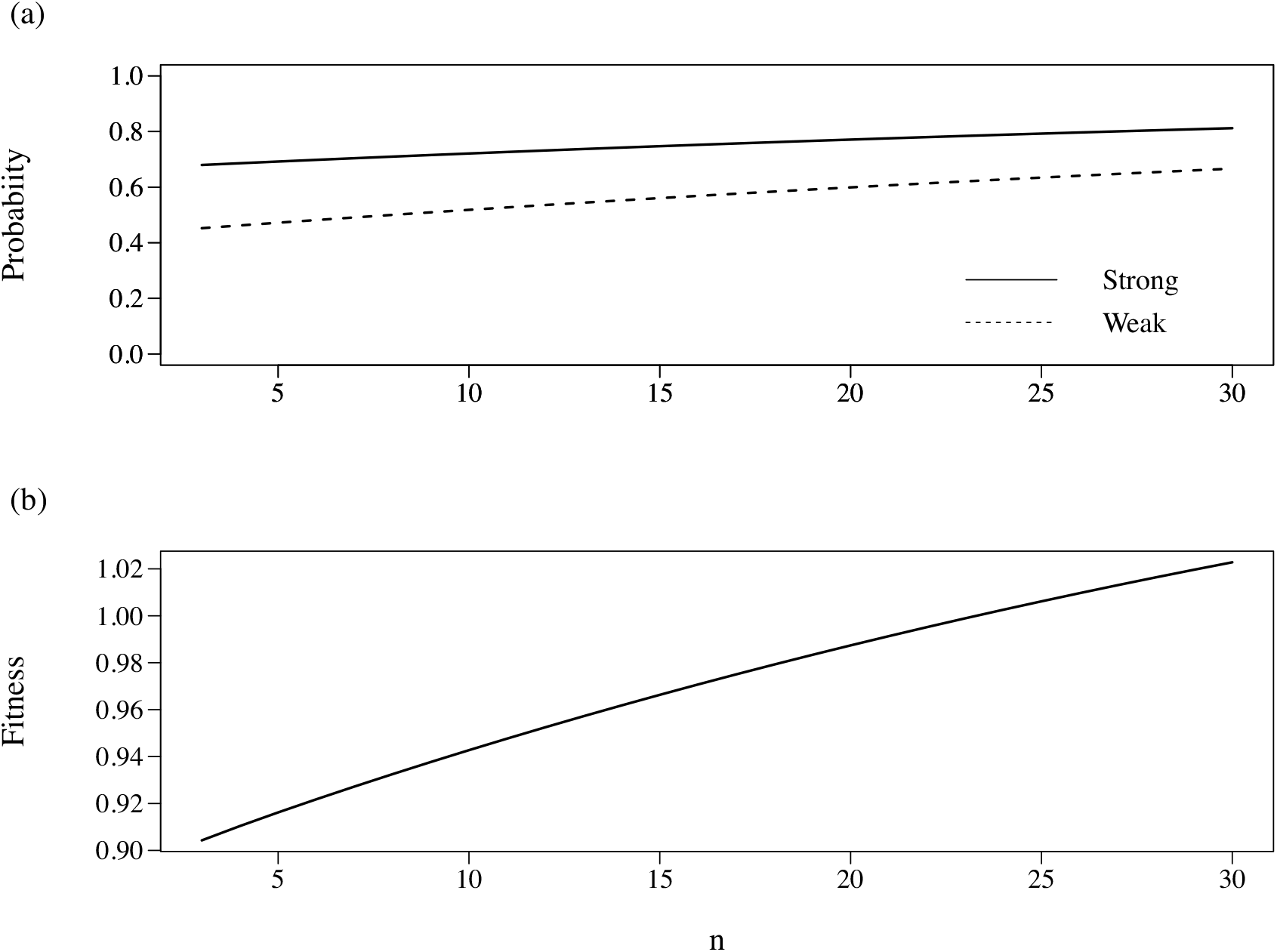
Probability of winning the coalition and fitness for various values of *n* at the initial condition. The initial frequencies of the four strategies are (*x*_00_,*x*_01_,*x*_10_,*x*_11_) = (0.97,0.01, 0.01, 0.01). We set *γ* =1, *s_s_* = 2, *s_w_* = 1, *R* =1, *α* = 2 and *c* = 1.5.

As *n* increases, the probability of winning the coalition and the fitness value of greedy individuals increases. This can be because larger groups can include a larger number of *C*_1_ individuals, resulting in higher probability of winning the coalition.

Figure 6 shows the effects of *ϕ* on the probability of winning the coalition and the fitness value of greedy individuals.

**Figure 6:**
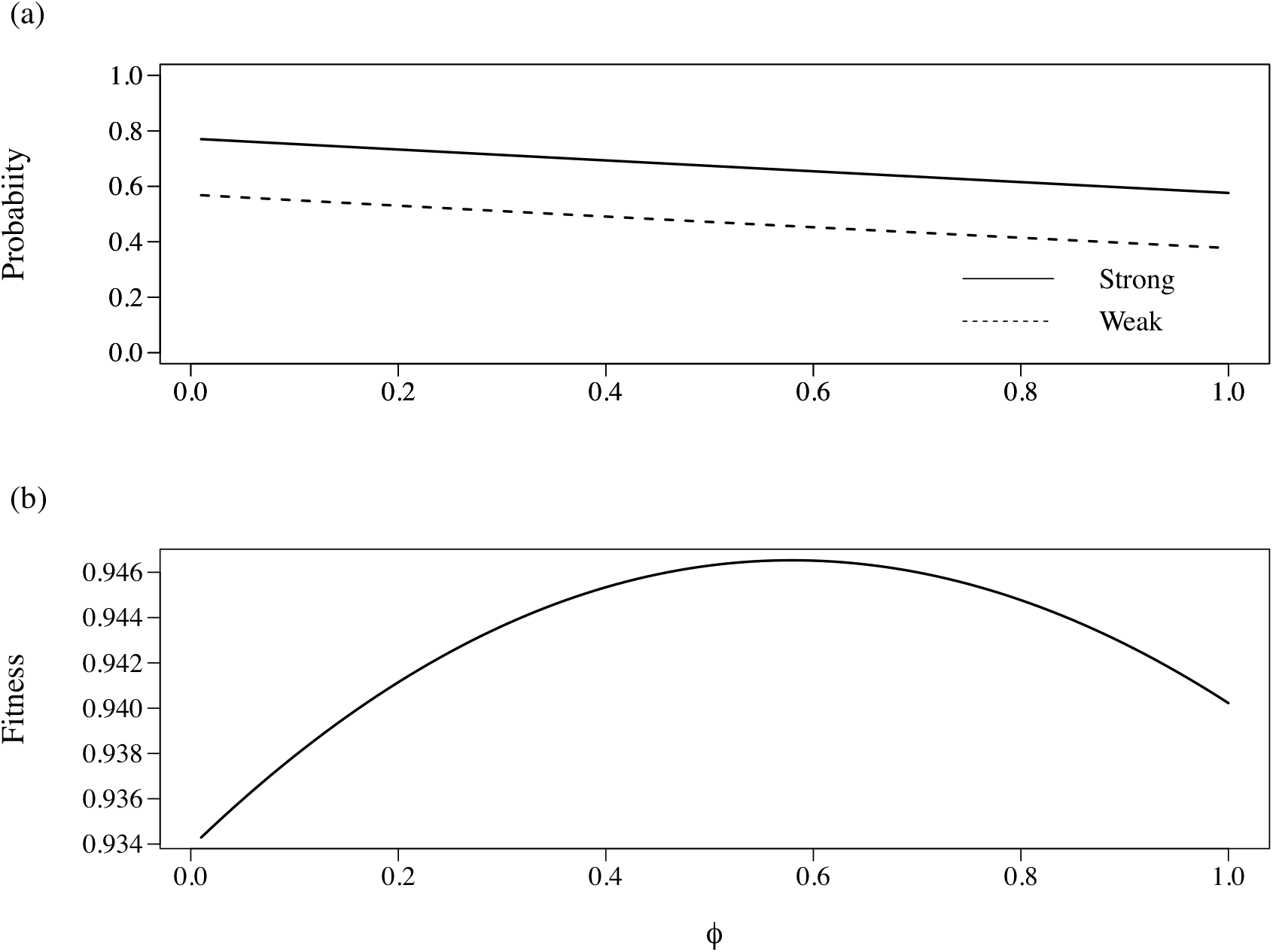
Probability of winning the coalition and fitness for various values of *ϕ* at the initial condition. The initial frequencies of the four strategies are (*x*_00_,*x*_01_,*x*_10_,*x*_11_) = (0.97,0.01, 0.01, 0.01). We set *γ* =1, *s_s_* = 2, *s_w_* = 1, *R* =1, *α* = 2 and *c* = 1.5.

As *ϕ* increases, the probability of winning the coalition decreases. The optimal value of *ϕ* exists to maximise the fitness value of greedy individuals.

## Discussion

In the present study, we investigated the evolution of egalitarian behaviour based on an n-player game extension of Gavrilets’s [24] model. Our results suggest that the evolution of egalitarian behaviour can be promoted by (i) potentially despotic payoff structure(*β*), (ii) large group size (*n*), (iii) small cost of competition (*c*) and (iv) variation in the strength of individuals (*ϕ*). In supplementary information, we also investigated effects of the synergy of the coalition, *α*, and the strength of a strong individual, *s_s_*, on the evolution of egalitarian behaviour. Our results suggest that effects of *α* is minor but large difference in fighting ability between strong and weak individuals can promote the evolution of egalitarian behaviour (Figure S1-S4). Gavrilets [24] reported that a potentially despotic payoff can promote the evolution of egalitarianism. Consistent with Gavrilets’s [24] results, our result also confirmed that a potentially despotic payoff structure can significantly affect the evolution of an egalitarian society.

The evolutionary transition from a population of monopoliser individuals to that of egalitarian individuals can be divided into two stages as follows. First, the frequency of greedy individuals increases if the fitness value of strong greedy individuals outperforms that of strong monopoliser individuals. Second, after the frequency of greedy individuals exceeds a certain level, greedy individuals are punished by a coalition. The frequency of egalitarian individuals, who joined the coalition and are not punished, increases, and eventually egalitarian individuals dominate the population. The four factors we mentioned above can contribute to the first stage. When the payoff structure is potentially despotic, that is, *β* is large, the fitness of the most successful type of individuals is representative. The fitness of each individual is composed of that of strong and weak individuals. As Figure 2 shows, the fitness values of strong greedy individuals are the largest but those of weak greedy individuals are less than those of monopoliser individuals. A large value of *β* emphasizes the fitness of strong greedy individuals and weakens the disadvantage of weak greedy individuals. As a result, a large value of *β* can be advantageous at the first stage.

When group size, *n*, is large, larger groups can include a larger number of *C*_1_ individuals, resulting in higher probability of winning the coalition. This effect is more important at the first stage than at the second stage, because there are a lot of *C*_1_ individuals at the second stage; that is, a group has a sufficient number of *C*_1_ individuals even if *n* is small.

Cost of competition, *c*, occurs only when *C*_1_ individuals exist. At the initial condition, because the majority of the population comprises monopoliser individuals, many monopoliser individuals interact with other monopoliser individuals. In this case, there is no competition. Therefore, only a small proportion of monopoliser individuals suffer the cost of competition, and small *c* is beneficial for greedy individuals or the evolution of egalitarian behaviour.

*ϕ* also works at the first stage and has an advantage and disadvantage for the evolution of greedy individuals. An advantage is an increase in strong individuals. An adaptive advantage of greedy individuals is through strong individuals defeating the owner: as shown in Figure 2, the fitness value of strong greedy individuals is larger than that of strong monopoliser individuals, while that of weak greedy individuals is less than that of weak monopoliser individuals. Therefore, increase in *ϕ* also increases strong individuals, which could provide an adaptive advantage to greedy individuals. A disadvantage is the decrease in the probability of winning the coalition. At the initial condition, since the frequency of *C*_1_ individuals is very low, the number of individuals joining the coalition is very small. As *ϕ* increases, the owner is more likely to be strong and thus less likely to be defeated; that is, the probability that the coalition wins decreases (Figure 6(a)). As a result, the optimal value of *ϕ* is determined based on the balance of the above-mentioned advantages and disadvantage.

In this study, we assumed the repeated interaction. In the supplementary information, we also examined the one-shot interaction. Egalitarian behaviour is more likely to evolve in the case of one-shot interaction than that of repeated interaction, while the qualitative tendencies are the same in both cases (Figures S5 and S6).

The results in the main text suggest that large group size can promote the evolution of egalitarian behaviour. In supplementary information, we investigated a model in which, when a coalition wins a competition over the resource, individuals who joined the coalition equally distribute the resource to all group members rather than share just within the coalition. In this situation, the evolution of egalitarian behaviour is likely to occur in a small group (Figures S8), which is inconsistent with the results in the main text. These discrepancies suggest that the heavy dependence of the evolution of sharing behaviour on group size is strongly related to the range of sharing.

In our model, egalitarian ethos has two components, resource sharing and punishment of monopolisers. Although a monopoliser is rarely observed in ethnographic records and thus punishment of monopolisers is also not likely to be recorded, our model implies that the evolution of egalitarian ethos is difficult to achieve by only pacifist individuals. Further, our results show that the evolution of egalitarian individuals occurred after the evolution of greedy individuals, implying that the evolution of resource sharing could follow the evolution of coalition formation.

Our model is based on Gavrilets’s [24] model. However, there are several differences. First, Gavrilets’s model is stochastic, while our model is deterministic. Second, Gavrilets’s model assumes (potentially) triadic interaction, while our model considers n-player interaction. Third, in Gavrilets’s model, the bully who attempts to take an amount of resource from the owner is a possible norm violator. On the other hand, in our model, the owner could be a monopoliser if he or she did not share the resource. Fourth, Gavrilets’s model tracks the evolution of the escalation threshold of each individual. If the difference in the strength of two individuals is smaller than the focal individual’s escalation threshold, the individual behaves in an aggressive manner. In our model, however, each individual’s decision making only depends on the owner’s behaviour.

Our model makes some unrealistic assumptions. First, we assume that individuals form a group in each generation. Second, we also assume that the strength of individuals is dichotomised into strong and weak. In reality, the strength of individuals should be distributed continuously. Third, actual individuals may consider the strength of opponents, although our model, as well as Gavrilets’s [24] model, neglects this factoIncorporating such factors can warrant further investigation.

